# Pharmacogenetic variants associated with off-target adverse drug reactions are mostly predicted to be benign

**DOI:** 10.1101/780213

**Authors:** Hannah McConnell, Matthew A Field, T. Daniel Andrews

## Abstract

Tools that predict the functional importance of genetic variation almost always rely on sequence conservation across deep evolutionary divergences as a primary discriminator. However, sequence conservation information is misleading when predicting the functional importance of pharmacogenetic variants related to off-target adverse drug reactions. Sequence conservation is largely maintained by evolutionary purifying selection, which has not been relevant for most drugs until very recently, especially for off-target effects. Here, we use a simple classification criteria to identify variants with off-target pharmacogenetic effects from the PharmGKB database. We show that off-target pharmacogenetic variation is predicted mostly to be benign by all state-of-the-art prediction tools we tested. Hence, off-target pharmacogenetic variants are overwhelmingly invisible to all predictive methodologies currently employed. Very different analytical approaches will be needed to address this important problem.

**Author Summary:** When a personal genome sequence is obtained for a given person, the sequence is compared to the human reference sequence to identify where it differs from the genome of that person. One application of this information is that it may identify how a specific person may react to particular drugs. However, when computationally predicting the functional importance of a genetic variant, the tools used rely heavily on sequence conservation information to make their prediction. From an evolutionary point of view, the use of drugs to treat diseases is a very recent activity – and one that has not had time to cause certain variants to either be selected for or removed from the population. This produces a blind-spot for tools that predict variant functional effects, especially for drugs with off-target interactions that may produce unanticipated effects.

## Introduction

Adverse drug reactions are historically classified into two main categories, which describe a general mechanistic distinction (1,2). Type A reactions are common and their effects are predictable and mostly dose-dependent. Type A reactions relate to interactions of a drug with its’ primary drug target. Conversely, Type B reactions are less common and are mostly unrelated to the main pharmacological action of the drug. Type B reactions, sometimes also called idiosyncratic drug reactions (3), can be dose-dependent or dose-independent, may be immunologically-mediated and/or may involve off-target drug interactions (2). Immune-mediated Type B reactions involve the drug inducing a specific immune response, such as the development of a skin rash commonly caused by administration of penicillin (4). Off-target drug effects can also occur without an immunological component, such as the interactions of anaesthetics with the ryanodine receptor 1 (RYR1) protein causing malignant hyperthermia (5).

Multiple databases aggregate, curate and annotate the growing body of identified pharmacogenetic variation (6,7). The Pharmacogenomic Knowledgebase (PharmGKB)(8) is the largest, open database of pharmacogenetic data, and at time of publication, includes information on nearly 150 pathways and over 20,000 individual variant annotations. Variants within PharmGKB are also annotated with effect types (dosage, efficacy, toxicity) and the level of confidence (categories 1-4) of the pharmacogenetic association, with category 1 being the highest. The pharmacogenetic variants included in PharmGKB cover a wide range of mutation types, from missense and synonymous variation to non-coding, intergenic and copy number variants.

The sheer number of genetic variants identified from a single, personal genome sequence is several orders of magnitude larger than the numbers of changes backed by experimentally-verified functional data. This interpretation gap is presently filled by a growing number of mutation function inference tools, including the well-known PolyPhen2 (9), CADD (10) and SIFT (11) tools. These inferential tools integrate sequence-based and, often, structural information to predict whether mutations and genetic variants are functionally different to a baseline reference sequence (12). In almost all cases, if a variant or mutation lies in a highly conserved region in a multispecies alignment of orthologous gene sequences, the variant will very likely be considered deleterious or damaging. Conversely, should the variant be broadly similar to sequence variation in this same alignment, the variant will be considered benign or functionally homologous. While some tools do consider structural information pertaining to the missense change, most of these tools also do incorporate sequence conservation data in their algorithms. Hence, the strength of purifying selection, removing non-functional or poorly functional variants from the population – and therefore ensuring conservation at the sequence level – is the strongest information presently used to classify a variant as either benign or deleterious.

The dependence of the algorithms on sequence conservation raises a clear question when assessing pharmacogenetic variation: has the variation been subject to purifying selection over evolutionary timescales? Much Type B pharmacogenetic variation will definitely not have been. Such pharmacogenetic variation violates a key assumption of the variant function inference tools and it will only be by chance that these tools correctly classify these variants.

A recent appraisal of mutation functional inference methods applied to pharmacogenetic missense variants found them to perform poorly (13). This effect was attributed to the ill-suited training sets used to build the models on which the algorithms rely, which prompted development of a better tool trained on pharmacogenetic variants present in just Absorption, Distribution, Metabolism and Excretion (ADME) process genes. ADME genes and the pharmacogenetic variants they contain are almost all of Type A, and analysing these separately from the off-target effects concentrated in Type B will definitely have strongly contributed to the improvement gains identified in this study.

We investigated pharmacogenetic variants held within the PharmGKB and sub-classified those that are Type B, off-target or idiosyncratic to determine whether the functional inferences made for these variants are different compared to pharmacogenetic variants in Type A genes. The PharmGKB contains substantial numbers of pharmacogenetic variants, across all variant evidence levels, that are computationally predicted to be benign. Within these benign variant groups, we show that these are overwhelmingly off-target and idiosyncratic variants.

## Results

### Distributions of pharmacogenetic variant functional inferences

Functional inference scores were obtained for 561 missense single nucleotide variants (SNVs) with Reference SNP (RS) cluster identifiers from the PharmGKB database using eleven different prediction tools (SIFT(11), PolyPhen2(14), CADD(10), DANN(15), FATHMM(16), GERP++(17), MutPred(18), Mutation Assessor(19), Mutation Taster(20), REVEL(21) and PhastCons(22)) (Supplementary Table S1). The distributions of scores from six of these (CADD, PolyPhen2, SIFT, MutationAssessor, MutPred and REVEL) are plotted in Figure 1. The predictions calculated for these functional variants ranged widely from benign to deleterious. For comparison, we also selected a random set of 2155 human missense SNVs with assigned RS cluster identifiers. The pattern of variant inferences is broadly similar across all tools and no single tool produces inferences that are qualitatively different (note that the scale of scores calculated by SIFT run in the opposite direction to other tools). These tools represent the range of methodologies available for mutation functional prediction and the categories of information used by each tool is annotated in Figure 1. Of interest, from this analysis the PharmGKB dataset includes many variants that are predicted to be functionally unimportant, even among variants with highest confidence. Of the 119 variants in category 1, the majority of predictions were deleterious (PolyPhen2 median score 0.996), though 6 variants were classed benign. The 183 variants in category 2 had a much broader range of predicted functional effects. The median score of variants in this category was benign (PolyPhen2 median score 0.138) was less than the median score of the randomly-chosen variants (PolyPhen2 median score 0.245) and 33 variants were predicted benign. The distribution of functional effect predictions in category 3 was strongly skewed towards benign variants (PolyPhen2 median score 0.012) and category 4 had a distribution very similar to the random variant set (PolyPhen2 median score 0.319). This pattern of inferred functional effects for PharmGKB variants was visually very similar between tools (Figure 1).

**Figure 1.**
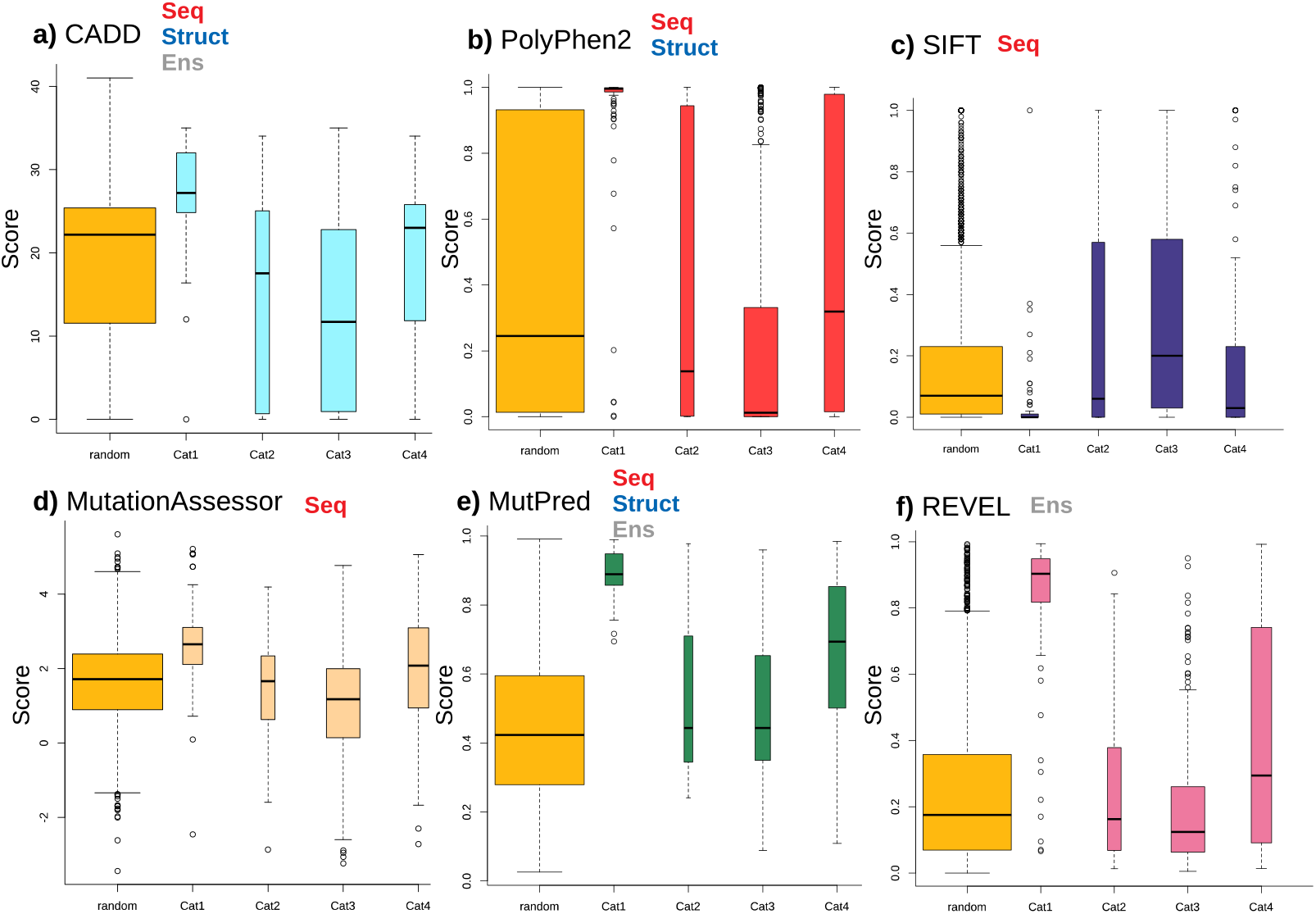
Distribution of functional effect scores of PharmGKB variants predicted by six mutation effect inference tools. Boxplots shown are of **a)** CADD Phred score, **b)** PolyPhen2 score, **c)** SIFT score, **d)** Mutation Assessor score, **e)** MutPred score and **f)** REVEL score. Scores are plotted for each tool in variant confidence categories (from 1 (highest) to 4 (lowest)) assigned by the PharmGKB annotation. Each tool is annotated with the information types it employs to make predictions – **Seq**: sequence conservation, **Struct**: protein structural metrics, **Ens**: an ensemble tool that integrates results of several individual tools.

A prior study (13) that noted functional prediction tools do not perform well on pharmacogenetic variation. Apart from the differing levels of evidence annotated to each PharmGKB variant, the known or suspected action of each variant is annotated into categories of Dosage, Efficacy, Toxicity/ADR and Other. Especially, the predicted-benign variants in PharmGKB are of note and may reflect the mechanism of these variants compared to the other variants in PharmGKB database. Specifically, we hypothesised that the majority of pharmacogenetic variants predicted to be of benign effect were of Type B pharmacogenetic mechanisms.

### Classification of pharmacogenetic variation to detect off-target effects

To investigate the possibility that Type B pharmacogenetic variants are predominantly predicted to be benign, we devised a simple classification system of PharmGKB variants (described in *Materials and Methods* and Table 2) that would be selective for those of Type B. From a possible 140 variants from categories 1A, 1B, 2A and 2B from PharmGKB, this classification system identified 24 variants as potentially Type B or off-target variants (Table 1). Half of these (12) were missense variants and the remainder being at non-coding or synonymous sites. Of the 12 missense variants, six were predicted as benign by PolyPhen2 or had a CADD phred score of less than 20 (a conservative threshold for deleterious variants). All variants, including non-coding variants were scored with CADD, of these only five had a phred score greater than 20. Hence, even of these high confidence pharmacogenetic variants, only a minority are predicted to be functionally important by existing tools.

**Table 1.** High confidence PharmGKB variants sorted with criteria to identify possible off-target effects.

| Variant | Gene | PharmGKB level of evidence | Example PharmGKB Chemicals | PharmGKB Phenotypes | CADD Phred score | PolyPhen category (score) | Mechanism Known? | Category | Variant type |
| --- | --- | --- | --- | --- | --- | --- | --- | --- | --- |
| rs1050828 | G6PD | 1B | chlorproguanil, dapsone | Malaria | - | Benign (0.202) | Yes (25) | Off-target | Missense |
| rs267606617 | MT-RNR1 | 1B | aminoglycoside antibacterials | Ototoxicity | - | Benign (0) | Yes (26) | Off-target | Non-coding |
| rs111888148 | RYR1 | 1A | desflurane, enflurane, halothane | Malignant Hyperthermia | 27.5 | Probably Damaging (0.998) | Limited (27) | Off-target | Missense |
| rs1800559 | CACNA1S | 1A | desflurane, enflurane, halothane | Malignant Hyperthermia | 24.4 | Probably Damaging (0.998) | Yes(28) | Off-target | Missense |
| rs1799971 | OPRM1 | 2B | ethanol | Alcoholism | 24 | Possibly Damaging (0.775) | Yes (29) | Off-target | Missense |
| rs1127354 | ITPA | 2B | peginterferon alpha-2b, ribavirin | Hepatitis C | 22.7 | - | Yes (30) | Off-target | Missense |
| rs1933437 | FLT3 | 2B | sunitinib | Carcinoma, Leukopenia | 21.4 | Possibly Damaging (0.69) |  |  | Missense |
| rs1801394 | MTRR | 2B | methotrexate | Lymphoma | 20.9 | Probably Damaging (0.99) | Proposed (31) | Off-target | Missense |
| rs2228001 | XPC | 1B | cisplatin | Neoplasms | 18.95 | Benign (0) | Yes (32) | Off-target | Missense |
| rs6025 | F5 | 2A | hormonal contraceptives | Thrombosis | 18.92 | Benign (0) | Yes (33) | Off-target | Missense |
| rs738409 | PNPLA3 | 2B | prednisolone, vincristine | Lymphoma | 15.7 | Possibly Damaging (0.906) | Yes (34) | Off-target | Missense |
| rs2232228 | HAS3 | 2B | anthracyclines | Cardiomyopathies | 15.61 | - | Proposed (35) | Off-target | Synonymous |
| rs16969968 | CHRNA5 | 2B | nicotine | Tobacco Use Disorder | 14.97 | Benign (0.011) | Yes (36) | Off-target | Missense |
| rs1800497 | ANKK1,DRD2 | 2B | ethanol | Alcoholism | 10.6 | Benign (0) | Yes (37) | Off-target | Missense |
| rs1051730 | CHRNA3 | 2B | nicotine |  | 10.1 | - | No |  | Synonymous |
| rs1076560 | DRD2 | 2B | cocaine | Cocaine-Related Disorders | 8.805 | - | No |  | Intronic |
| rs746647 | CCHCR1 | 2B | nevirapine | HIV | 5.575 | - | No |  | Intronic |
| rs4693075 | COQ2 | 2B | statins | Muscular Diseases | 2.092 | - | No |  | Intronic |
| rs10497203 | TANC1 | 2B | radiotherapy | Prostatic Neoplasms | 1.974 | - | Proposed (38) | Off-target | Intronic |
| rs730012 | LTC4S | 2B | aspirin | Urticaria | 1.936 | - | No |  | 5'-flanking |
| rs1872328 | ACYP2 | 2B | cisplatin | Brain Neoplasms, Ototoxicity | 0.897 | - | Limited (39) | Off-target | Intronic |
| rs716274 | DYNC2H1 | 2B | etoposide, platinum compounds |  | 0.621 | - | No |  | Intergenic |
| rs489693 | MC4R | 2B | clozapine, risperidone | Autism, Schizophrenia | 0.325 | - | No |  | Intergenic |
| rs7779029 | SEMA3C | 2B | irinotecan | Carcinoma | 0.165 | - | No |  | Intronic |
| rs1517114 | C8orf34 | 2B | irinotecan | Carcinoma | 0.081 | - | No |  | Intronic |

**Table 2.** Classification criteria used to identify off-target pharmacogenetic variants from the PharmGKB database.

| Step | Filter |
| --- | --- |
| 1 | Variant only has type:toxicity/ADR |
| 2 | For drug and gene pairs, there are no other variants with effect type other than 'Toxicity/ADR' |
| 3 | Gene containing variant is NOT an ADME process gene OR annotated in GO with 'xenobiotic metab process' OR 'transporter') |
| 4 | Exclude synonymous and non-coding variants |
| 5 | Literature review indicates a known mechanism, which is an off-target drug/protein interaction |

For each variant in Table 1, a manual literature survey was conducted to further classify each variant by mechanism – and whether these were off-target, Type B pharmacogenetic variants. Of the 24 variants, the mechanism of action was known in 15 cases and every one of these had an off-target (or Type B) effect, including four cases where the variant was not a missense variant. This included all five high-confidence category 1 pharmacogenetic variants. Treating this result as strong validation of the classification system we applied to identify off-target effects, we applied this scheme to lower evidence variants from PharmGKB categories 3 and 4.

Given that category 3 variants were strongly skewed towards being predicted as benign (Figure 1), we asked the question whether this category of variants is overwhelmingly off-target, Type B pharmacogenetic variants. Where the mechanism of pharmacogenomic effect was known, 12 out of a total 17 variants we manually appraised were off-target effects (Supplementary Table S2). This result provides a measure of the efficacy 70% (12/17) of this method to detect off-target pharmacogenetic variants purely from data analysis without associated mechanistic studies. Further precision may be obtained through excluding potential off-target variants with a deleterious functional prediction.

### Discussion

In this work we have classified pharmacogenetic variants by their mechanism, where it is known, to appraise whether off-target pharmacogenetic variants are consistently predicted to be less deleterious than other functionally-important variants. In predicting the functional importance of a missense variant, the current generation of mutation inference tools make distinctions on the basis of sequence conservation – where a variant occurring in a conserved region are highly likely to be classified as deleterious. Conservation in sequence data between orthologous sequences implies that during evolution, the function of the encoded protein relied on there being either no changes or else only conservative changes. The action of drugs prescribed to human patients is a very recent occurrence in an evolutionary context and unless a pharmacogenetic variant is related to the normal function of cellular processes, such as ADME processes, there will be no information present in the sequence record from which to detect functional importance. Off-target and/or idiosyncratic pharmacogenetic variants have been too recent a selective condition for evolutionary processes to have left their mark at the nucleotide sequence level, even within the human genome. Hence, when looking to identify the potential functional importance of a pharmacogenetic variants, these will be mostly invisible to current inferential tools. Our results show that in the category of pharmacogenetic variants that have an off-target mechanism, these are almost completely classified as benign, predicted to be of no functional importance.

Given the reliance of variant functional inference tools on sequence conservation, it is unsurprising that the tools perform poorly on pharmacogenetic variation that have not been subjected to purifying selection. Without this sequence conservation information however, what methods and datasets remain available to differentiate between benign and functionally important variation? Tools that use protein structural information are intuitively a good fall-back, yet our investigation showed little difference between tools, even those that make use of structural features to predict deleterious variation. Potentially, tools to have no reliance on sequence conservation will be needed to test this possibility rigorously.

Pharmacogenetic missense variants represent a complex set of genetic factors with highly diverse functional mechanisms by which they influence drug efficacy. Functional predictions of the likely impact of a given variant are driven by measures of sequence conservation over deep evolutionary timescales. Unlike pathogenic variants identified in rare genetic diseases, the action of drugs has not necessarily been subject to evolutionary selective forces and purifying selection – and, at least, is applied to an as-yet very limited range of species. Many pharmacogenes containing variants with a Type A mechanism will have been subject to purifying selection, as the drug is most often just another xenobiotic compound which the target-protein acts on. However, drugs that cause a Type B or off-target effect are much less likely to be subject to the same selection. This mechanistic difference is important when sequence conservation information is pivotal to the prediction of the functional effect of a sequence variant. We show most off-target pharmacogenetic variants of this type are predicted to be functionally unimportant. New methods that can detect pharmacogenetic variation that has not been subjected to purifying selection are needed.

## Materials and Methods

### Pharmacogenetic Variant Datasets

A set of pharmacogenetic variants with Reference SNP (RS) cluster identifiers were obtained from PharmGKB (8) with custom code to combine variant annotations. Variants within PharmGKB are classified by gene, type of effect, level of evidence, specific drug, chemical, disease and phenotype. Variants were further annotated with Variant Effect Predictor(23).

### Classification of Off-Target Pharmacogenetic Variants

A simple classification scheme was devised to identify and confirm likely off-target variants (Table 1). All clinical variants with evidence category of 1A, 1B, 2A and 2B from the PharmGKB database (8) were first filtered for variants with PharmGKB annotation of effect type of ‘Toxicity/ADR’ for any particular chemical (drug). Variants were removed if they also had an additional effect type (other than ‘Toxicity/ADR’) for the same drug. If different drugs produced different effect types for the same variant, these were not considered as criteria for variant filtering. Next, variants were removed if they were present in ADME process genes (categorized as such in the PharmaADME database; www.pharmaadme.org) or were annotated with Gene Ontology (24) categories of ‘xenobiotic metabolism process’ or with ‘transporter’. Of these remaining variants, synonymous and non-coding variants were excluded, leaving just missense variants. With this filtered list, the cited literature for each variant was appraised to discern whether the variant was an off-target effect. Variants were retained where a molecular mechanism for the adverse drug effect was known, did not involve a protein or cellular system related to the intended drug effects or did not likely alter the protein abundance of the intended drug targets.

### Functional Effect Prediction

The predicted functional effect of mutations was predicted with SIFT(11), PolyPhen2(14), CADD(10), DANN(15), FATHMM(16), GERP++(17), MutPred(18), Mutation Assessor(19), Mutation Taster(20), REVEL(21) and PhastCons(22), relative to EnsEMBL canonical transcripts with the Variant Effect Predictor (VEP)(23).

## Acknowledgements

The authors would like to thank the National Computational Infrastructure for continued access to computational resources. This work has been partially supported by NHMRC fellowship APP1139756 to MAF.

## Author Contributions

TDA, MAF and HM devised the research. TDA, MAF and HM performed the analysis. TDA, MAF and HM wrote the paper.

